# Opioid receptors reveal a discrete cellular mechanism of endosomal G protein activation

**DOI:** 10.1101/2024.10.07.617095

**Authors:** Nicole M. Fisher, Mark von Zastrow

## Abstract

Many GPCRs initiate a second phase of G protein-mediated signaling from endosomes, which inherently requires an increase in G protein activity on the endosome surface. G_s_-coupled GPCRs are thought to achieve this by internalizing and allosterically activating cognate G proteins again on the endosome membrane. Here we demonstrate that the μ-opioid receptor (MOR), a G_i_-coupled GPCR, increases endosomal G protein activity in a different way. Leveraging conformational biosensors, we resolve the subcellular activation dynamics of endogenously expressed MOR and G_i/o_-subclass G proteins. We show that MOR activation triggers a transient increase of active-state G_i/o_ on the plasma membrane that is followed by a prolonged increase on endosomes. Contrary to the G_s_-coupled GPCR paradigm, however, we show that the MOR-induced increase of active-state G_i/o_ on endosomes requires neither internalization of MOR nor activation of MOR in the endosome membrane. We propose a distinct and additional cellular mechanism for GPCR-triggered elevation of G protein activity on endosomes that is mediated by regulated trafficking of the activated G protein rather than its activating GPCR.

## Introduction

G protein-coupled receptors (GPCRs) comprise the largest family of signaling receptors and integral membrane proteins in animals. In response to a biologically relevant stimulus, GPCRs increase the activity of heterotrimeric GTP-binding proteins (G proteins), which act as transducers to relay biological information downstream of the activated receptor. Individual GPCRs differentially elevate the activity of homologous G protein subclasses (G_s/olf_, G_i/o_, G_q/11,_ and G_12/13_), and these differ, in turn, in their functional regulation of downstream effector systems in the cell (1, 2).

Physiological signaling through this biochemical framework is now recognized to be organized across multiple subcellular compartments (3, 4). In particular, GPCR signaling initiated from endosomes has been linked to distinct outcomes compared to signaling initiated from the plasma membrane, indicating that compartmentalized signaling can impact the downstream cellular consequences of ligand-induced GPCR activation, reviewed in (5, 6). This fundamentally requires an activated GPCR to trigger an increase in G protein activity at the appropriate membrane location. G proteins are activated on the plasma membrane through an allosteric coupling reaction that requires the membrane-tethered G protein to transiently contact its activating GPCR embedded in the membrane bilayer (7, 8). While this activation mechanism has been well described at the plasma membrane for all G protein subclasses, it is less clear how G protein activity is increased on the endosome membrane. The present understanding has been developed largely through the study of G_s_-coupled GPCRs. G_s_-coupled GPCRs are thought to increase G protein activity on endosomes by internalizing and mediating a second round of allosteric coupling to cognate G proteins on the endosome limiting membrane. This model is supported by experiments that demonstrate that endosomal G protein-mediated signaling requires receptor internalization (9–15) and that its strength depends on the degree or duration of receptor activation in the endosome limiting membrane (16–20).

Here we show that endosomal G protein activation mediated by the μ-opioid receptor (MOR), a G_i_-coupled GPCR, is not compatible with this paradigm. Our results reveal the existence of a distinct and additional cellular mechanism for increasing cognate G protein activity on endosomes that does not depend on receptor internalization or activation in the endosome membrane. Instead, our results reveal a cellular mechanism of endosomal signaling that is mediated by trafficking of activated G proteins rather than their activating GPCRs.

## Results

### Endosomal localization of G_i/o_ increases following activation by MOR

As a first step toward delineating the spatiotemporal organization of G protein activation by MOR, we examined the subcellular localization of fluorescently-labeled G_i/o_-subclass isoforms by live-cell confocal microscopy. mNeonGreen was inserted into the αb-αc loop of G_i1_α, a tagging strategy previously demonstrated to preserve protein function (21). G_i1_-mNeonGreen, an endosome marker (FYVE-mApple), and a FLAG-tagged MOR construct (SSF-MOR) were co-expressed in HEK293 cells in which all endogenous G_i/o_ α-subunit genes (GNAI1, GNAI2, GNAI3, GNAO, GNAZ and GNAT) were knocked out using CRISPR-Cas9 (Gi-KO cells) (22). Receptors present in the plasma membrane were labeled by preincubation with an anti-FLAG antibody conjugated to AlexaFluor 647 (M1-647). In cells not exposed to opioid agonist, G_i1_-mNeonGreen localized primarily to the plasma membrane but was also observed on internal puncta, many of which were identified as endosomes by colocalization with FYVE-mApple (Figure 1A). Upon application of the opioid full agonist DAMGO we observed a qualitative increase in G_i1_-mNeonGreen fluorescence in SSF-MOR-containing endosomes (Figure 1A). Similar results were observed using mNeonGreen-labeled versions of G_i2,_ Gi_3_, G_oA,_ G_oB,_ and G_z_ (Figure S1). To confirm this increase and quantify relocalization to endosomes across a large number of cells, we used a luciferase protein complementation assay (NanoBiT). In this assay, MOR or G_i1_ were fused to LgBit (MOR-LgBiT or G_i1_-LgBiT), and accumulation on endosomes was measured by protein complementation with an endofin derived FYVE domain fused to SmBiT (FYVE-mApple-SmBiT) that localizes on the endosome limiting membrane. Application of DAMGO promoted MOR accumulation in endosomes (Figure 1B) with the expected time course based on receptor internalization measured previously using other methods (23). DAMGO also promoted G_i1_-LgBiT accumulation on endosomes (Figure 1C), consistent with the microscopy results (Figure 1A). Interestingly, this assay revealed that G_i1_-LgBiT accumulation on endosomes occurs with distinct and generally faster kinetics than endosomal accumulation of MOR-LgBiT.

**Figure 1.**
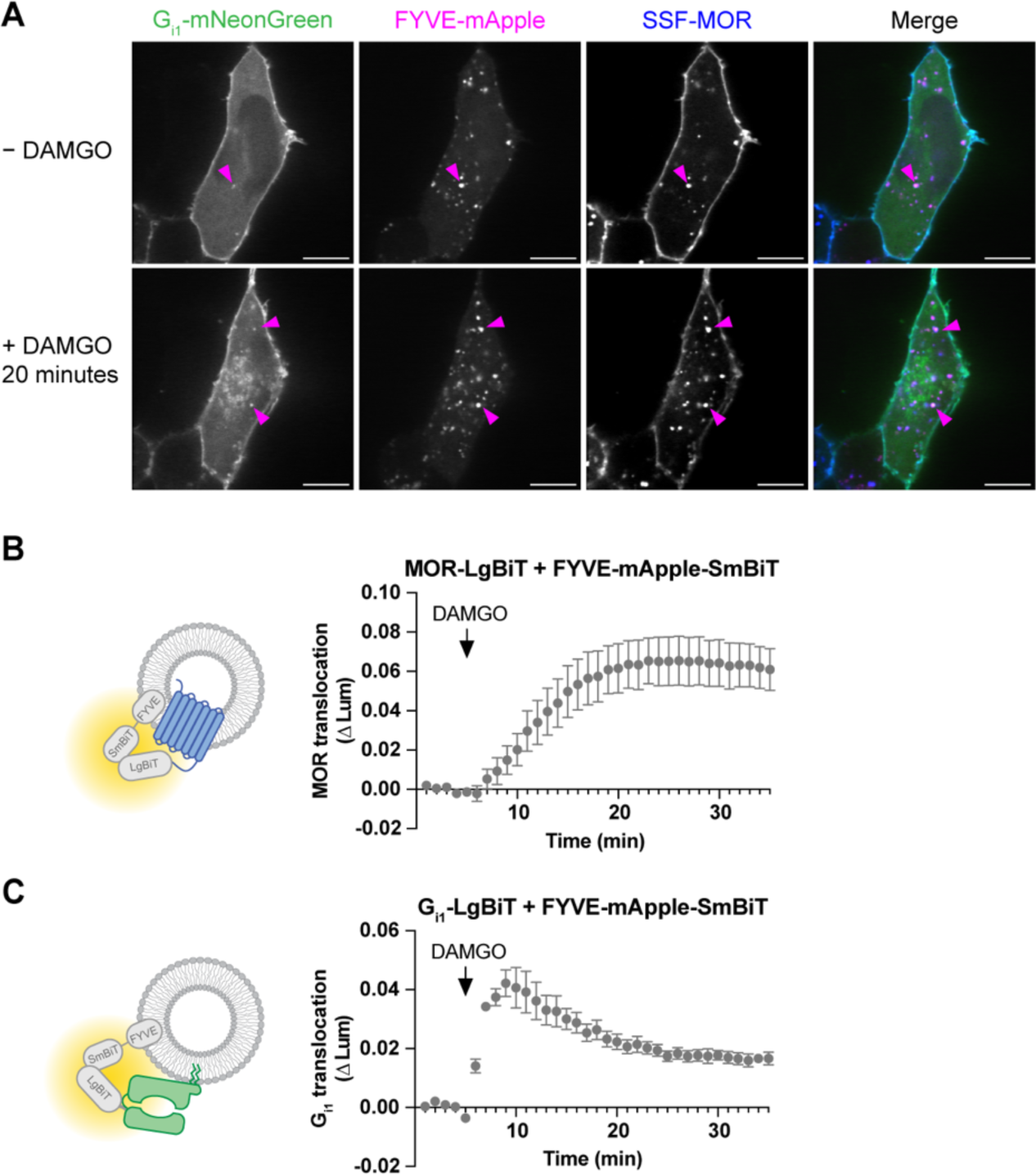
G_i1_ relocalizes to endosomes upon activation by MOR. A) Live imaging by confocal microscopy of Gi-KO HEK293A cells expressing G_i1_-mNeonGreen, the endosome marker FYVE-mApple and SSF-MOR labeled with ⍺-FLAG M1-647. Images depict frames before and after 20-minute treatment with 10 µM DAMGO. Magenta arrows indicate endosomes positive for G_i1_ - mNeonGreen. Representative images from 3 independent experiments. Scale bar = 10 µm. B) NanoBiT luminescence signal over time for cells expressing SSF-MOR-LgBiT + FYVE-mApple-SmBiT upon addition of 1 µM DAMGO. C) NanoBiT luminescence signal over time for cells expressing G_i1_-LgBiT + FYVE-mApple-SmBiT upon addition of 1 µM DAMGO. For NanoBiT assays, N=3 independent experiments. Error bars represent ± SEM.

### MOR and G_i/o_ are sequentially activated on the plasma membrane followed by endosomes

Having observed endosomal accumulation of G_i/o_ triggered by agonist-induced activation of MOR, we next tested for the presence of active-state MOR and G_i/o_ at each membrane location. To detect active-state MOR, we utilized a receptor-binding nanobody (Nb33) validated previously as an active-state biosensor by fluorescence microscopy (24) and adapted it for NanoBiT assays by fusion to SmBiT (Figure 2A). We measured recruitment of Nb33-SmBiT to LgBiT tethered to the plasma membrane (LgBiT-CAAX) or to the endosome membrane (FYVE-LgBiT). In HEK293 cells expressing SSF-MOR, DAMGO promoted sequential phases of Nb33-SmBiT recruitment, first to the plasma membrane and then to endosomes (Figure 2A-B), with a similar dependence on agonist concentration at each location (Figure 2C). To detect active-state G_i/o_, we chose an engineered G_i/o_ effector domain derived from RAP1GAP that was shown previously to specifically detect active-state G_i/o_ in intact cells (Gi-ED, (25)), and we similarly adapted this protein for the NanoBiT assay. Gi-ED-SmBiT detected sequential DAMGO-induced phases of endogenous G_i/o_ activation on the plasma membrane followed by endosomes. Verifying specificity for G_i/o_-subclass proteins, both signals were blocked by pretreatment of cells with pertussis toxin (Figure 2D-E, orange points). Each phase of DAMGO-induced G_i/o_ activation had a similar dependence on agonist concentration at each location (Figure 2F) and the concentration-dependence of G_i/o_ activation was left-shifted relative to MOR activation at both locations as expected (Figure 2C,F). Further verifying the location-specificity of biosensor detection, no recruitment signal was observed using either sensor in control experiments in which LgBiT was targeted to the mitochondrial outer membrane (Figure S2).

**Figure 2.**
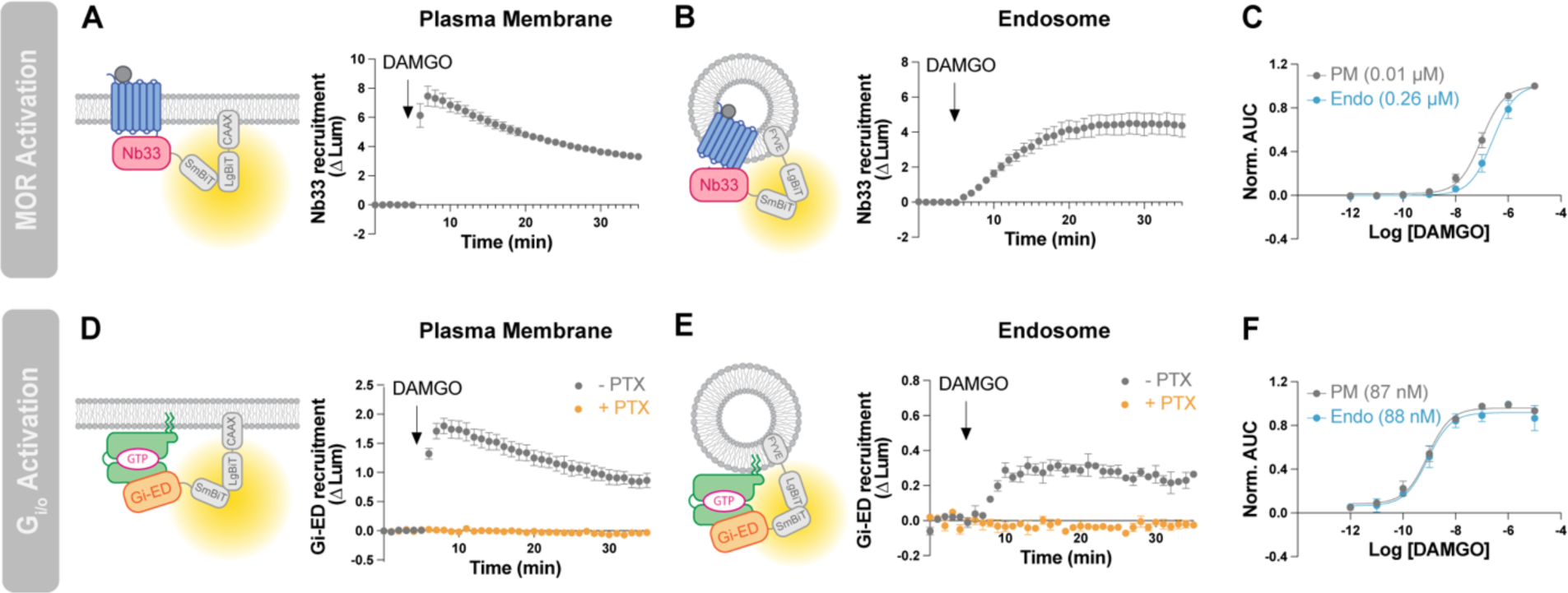
Sequential phases of MOR and G_i/o_ activation on the plasma membrane and endosomes. A) Nb33-SmBiT recruitment to the plasma membrane labeled with LgBiT-CAAX following addition of 1 µM DAMGO. B) Nb33-SmBiT recruitment to endosomes labeled with FYVE-LgBiT following addition of 1 µM DAMGO. C) Concentration response curves for Nb33-SmBit recruitment to each LgBiT construct. EC_50_ values noted in parentheses. D) Gi-ED-SmBiT recruitment to the plasma membrane labeled with LgBiT-CAAX following addition of 1 µM DAMGO. E) Gi-ED-SmBiT recruitment to endosomes labeled with FYVE-LgBiT following addition of 1 µM DAMGO. In panels D and E, orange curves represent cells pretreated with 100 ng/mL pertussis toxin (PTX). F) Concentration response curves for Gi-ED-SmBiT recruitment to each LgBiT construct. EC_50_ values are noted in parentheses. N=3 independent experiments. Error bars represent ± SEM.

We additionally confirmed specificity of Gi-ED-SmBiT in detecting activation of endogenous G_i/o_ genetically by demonstrating that DAMGO-induced activation of SSF-MOR failed to produce any detectable recruitment at either the plasma membrane or endosomes in Gi-KO cells (Figure S3, mock condition). Moreover, the recruitment signal was rescued at both locations by reexpression of the G_i/o_ constructs used in Figure 1 and Figure S1. Gi-ED-SmBiT detected activation at both the plasma membrane and endosomes in Gi-KO cells upon rescue with G_i1_, G_i2_, G_i3_, G_oA_ or G_oB_. The only exception was G_z_, where we failed to observe rescue at either location (Figure S3). Together, these data indicate that the observed sequential elevation of active-state G_i/o_ on the plasma membrane followed by endosomes is specific and widespread across G_i/o_ isoforms.

We also noted that the increase of active-conformation G_i/o_ level on endosomes, detected by Gi-ED-SmBiT, occurred more rapidly than the accumulation of active-conformation MOR detected by Nb33-SmBiT (T_max_= 23 min for Nb33-SmBiT, T_max_= 8 min for Gi-ED-SmBiT, Figure 2C,F). This observation, together with the generally faster kinetics of G protein trafficking to endosomes relative to MOR (Figure 1), motivated us to investigate the ligand-dependent regulation of endosomal G protein activity in more detail.

### Endosomal G_i/o_ activity does not depend on MOR activation in the endosome membrane

We began by examining the reversibility of MOR and G_i/o_ activation. We activated MOR by bath application of a moderate concentration of DAMGO (100 nM) and then reversed this by applying antagonist in excess (10 µM), using either naloxone, a membrane-permeant alkaloid antagonist, or CTOP, an antagonist peptide that is membrane-impermeant. Both antagonists fully reversed MOR activation at the plasma membrane, detected by Nb33-SmBiT, within 1 minute (Figure 3A), the temporal resolution limit of our assay, and the effects of each antagonist were indistinguishable in an area under the curve (AUC) analysis (Figure 3B). Naloxone also rapidly reversed the DAMGO-induced increase of active-state MOR in endosomes but CTOP, consistent with its membrane-impermeant nature, reversed the endosomal activation signal much more slowly (Figure 3C). This resulted in a significant difference when assessed by AUC analysis (Figure 3D). The slower time course of CTOP-induced loss of active-state MOR in endosomes (t_½_ ≥ 5 min) is consistent with the kinetics of MOR recycling (26), so we interpret the slow decrease in NanoBiT signal produced by CTOP as a result of receptor exit from endosomes for the recycling pathway.

**Figure 3.**
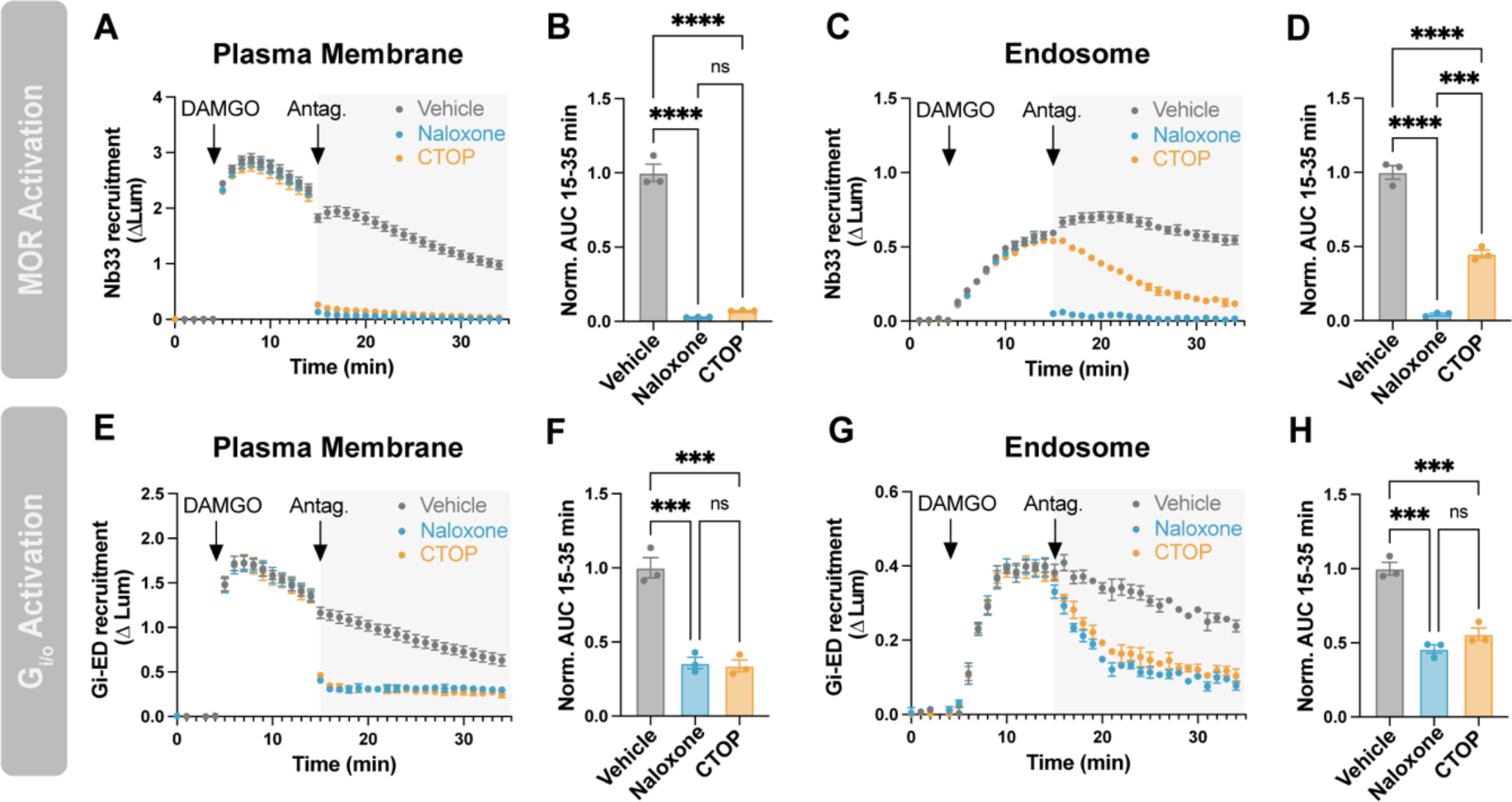
Endosomal G_i/o_ activation does not depend on active MOR in the endosome membrane. A) Nb33-SmBiT recruitment to the plasma membrane (LgBiT-CAAX) with addition of 100 nM DAMGO at 5 minutes followed by addition of 10 µM antagonist (naloxone or CTOP) at 15 minutes. B) Quantification of data in panel A. AUC was calculated from data collected after the antagonist add (15-35 min). C) Nb33-SmBiT recruitment to endosomes (FYVE-LgBiT) with addition of 100 nM DAMGO at 5 minutes followed by addition of 10 µM antagonist at 15 minutes. D) Quantification of data in panel C. AUC was calculated from data collected after the antagonist add (15-35 min). E) Gi-ED-SmBiT recruitment to the plasma membrane (LgBiT-CAAX) with addition of 100 nM DAMGO at 5 minutes followed by addition of 10 µM antagonist at 15 minutes. F) Quantification of data in panel E. AUC was calculated from data collected after the antagonist add (15-35 min). G) Gi-ED-SmBiT recruitment to endosomes (FYVE-LgBiT) with addition of 100 nM DAMGO at 5 minutes followed by addition of 10 µM antagonist at 15 minutes. H) Quantification of data in panel G. AUC was calculated from data collected after the antagonist add (15-35 min). One-way ANOVA with Tukey’s multiple comparisons, ***p<0.001, ****p<0.0001. N=3 independent experiments. Error bars represent ± SEM.

Both antagonists also rapidly reversed the DAMGO-induced increase of active-state G_i/o_ detected by Gi-ED-SmBiT at the plasma membrane (Figure 3E-F). Based on the present model of endosomal G protein activation, and the results described above, we expected naloxone to reverse the DAMGO-induced increase of active-state G_i/o_ on endosomes more quickly than CTOP. Surprisingly, this was not the case. Rather, both naloxone and CTOP reversed the G_i/o_ activation signal on endosomes with an indistinguishable time course (Figure 3G), and no difference was detected by AUC analysis (Figure 3H). A simple interpretation of these results is that the MOR-mediated elevation of active-state G_i/o_ on endosomes does not depend on the presence of active-state MOR in the endosome membrane.

### Endosomal activation of Gi/o does not require MOR internalization

If this is the case, we predicted that the MOR-triggered production of active-conformation G_i/o_ on endosomes should also be insensitive to experimental manipulations that suppress MOR internalization. We tested this using compound 101, a chemical inhibitor of GRK2/3 activity that strongly suppresses regulated endocytosis of MOR (27). As expected, compound 101 significantly suppressed the DAMGO-induced accumulation of active-state MOR in endosomes detected by Nb33-SmBiT (Figure 4C-D). Importantly, in contrast, compound 101 had no detectable effect on the elevation of active-state G_i/o_ on endosomes detected by Gi-ED-SmBiT (Figure 4G,H).

**Figure 4.**
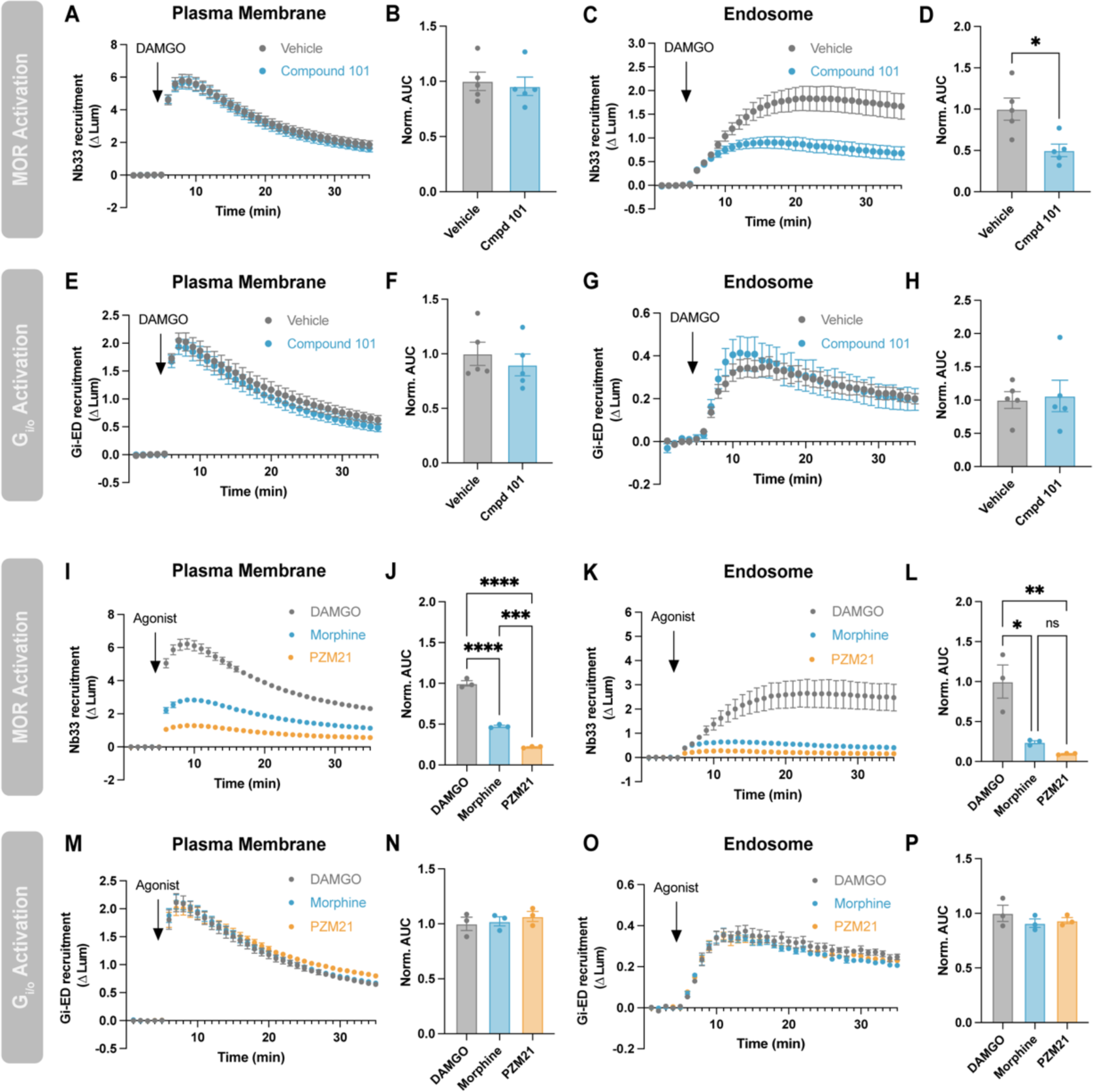
Endosomal G_i/o_ activation does not require MOR internalization. A) Nb33-SmBiT recruitment to the plasma membrane (LgBiT-CAAX) or C) to endosomes (FYVE-LgBiT) by 1 µM DAMGO with and without 20-minute pretreatment with 30 µM Compound 101. B,D) Quantification of AUC in A and C, respectively. E) Gi-ED-SmBiT recruitment to the plasma membrane (LgBiT-CAAX) or G) to endosomes (FYVE-LgBiT) by 1 µM DAMGO with and without pretreatment with 30 µM Compound 101. F,H) Quantification of AUC in E and G, respectively. I) Nb33-SmBiT recruitment to the plasma membrane (LgBiT-CAAX) or K) to endosomes (FYVE-LgBiT) by 1 µM DAMGO, morphine or PZM21. J,L) Quantification of AUC in I and K, respectively. M) Gi-ED-SmBiT recruitment to the plasma membrane (LgBiT-CAAX) or O) to endosomes (FYVE-LgBiT) by 1 µM DAMGO, morphine or PZM21. N,P) Quantification of AUC in M and O, respectively. One-way ANOVA with Tukey’s multiple comparisons, *p<0.05, **p<0.01, ***p<0.001, ****p<0.0001. N=3-5 independent experiments. Error bars represent ± SEM.

To test this hypothesis in a different way, we compared the effects of three chemically distinct agonists that differ in their ability to promote MOR internalization: DAMGO, a full agonist that robustly internalizes MOR; morphine, a partial agonist that weakly internalizes MOR (28); and PZM21, a less efficacious and G protein-biased partial agonist with minimal endocytic activity (29). The reduced agonist efficacies of morphine and PZM21 were reflected in differences in the level of Nb33-SmBiT recruitment that they produce on the plasma membrane at saturating concentration (Figure 4I-J). Also as expected, based on partial agonists stimulating MOR internalization poorly, morphine and PZM21 produced much less accumulation of active-state MOR in endosomes than DAMGO (Figure 4K-L). Remarkably, each agonist elevated active-state G_i/o_ on the plasma membrane and endosomes to comparable levels (Figure 4M-P). This suggested to us that MOR internalization is not required for G_i/o_ activation on endosomes.

### Endogenous MOR triggers endosomal G_i/o_ activation specifically from the plasma membrane

An alternative explanation for the above observations is that the expression level of recombinant MOR used in HEK293 cells produced such a high level of receptor reserve that effects of MOR internalization were simply missed. Therefore, we next sought to assess the role of MOR internalization in endosomal G_i/o_ activation in a cell model expressing endogenous receptors at a low level.

We focused on SH-SY5Y neuroblastoma cells because this neuronally derived cell line expresses endogenous MOR in much lower amount (∼100 fmol/mg (30)) than achieved by recombinant expression in HEK293 cells (typically > 1 pmol/mg) (31, 32). We found both Nb33-SmBiT and Gi-ED-SmBiT sufficiently sensitive to detect sequential phases of endogenous receptor and G protein activation, respectively, in this native cell model (Figure 5). Compound 101 did not prevent MOR activation in the plasma membrane, detected by Nb33-SmBiT recruitment, but it blocked the accumulation of active-state receptors in endosomes (Figure 5A-D). Further, compound 101 increased and prolonged the endogenous MOR-mediated activation of G proteins on the plasma membrane, detected by Gi-ED-SmBiT, consistent with compound 101 suppressing both MOR desensitization and internalization (Figure 5E-F). Remarkably, despite these effects, compound 101 actually *enhanced* the MOR-induced increase of active-state G protein accumulation on endosomes (Figure 5G-H). These data provide additional support for the interpretation that G_i/o_ activation at endosomes does not depend on coupling to activated MOR in endosomes. Rather, they suggest that the second phase of endosomal G_i/o_ activation is initiated by MOR activation of G_i/o_ on the plasma membrane.

**Figure 5.**
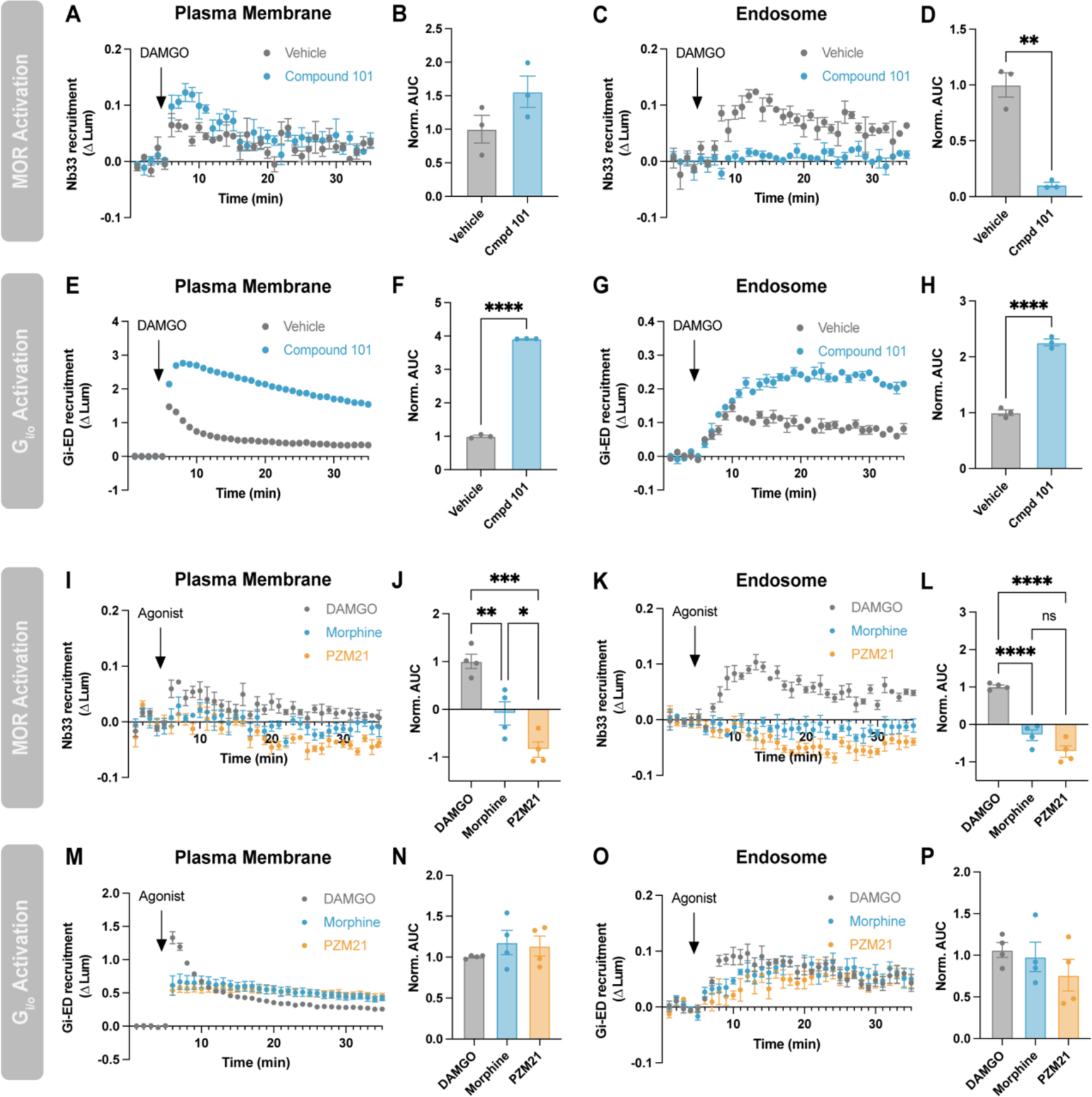
Surface MORs drive G_i/o_ activity on endosomes in SH-SY5Y cells. A) Nb33-SmBiT recruitment to the plasma membrane (LgBiT-CAAX) or C) to endosomes (FYVE-LgBiT) by 1 µM DAMGO with and without pretreatment with 30 µM compound 101. B,D) Quantification of AUC in A and C, respectively. E) Gi-ED-SmBiT recruitment to the plasma membrane (LgBiT-CAAX) or G) to endosomes (FYVE-LgBiT) by 1 µM DAMGO with and without pretreatment with 30 µM compound 101. F,H) Quantification of AUC in E and G, respectively. I) Nb33-SmBiT recruitment to the plasma membrane (LgBiT-CAAX) or K) to endosomes (FYVE-LgBiT) by 1 µM DAMGO, morphine or PZM21. J,L) Quantification of AUC in I and K, respectively. M) Gi-ED-SmBiT recruitment to the plasma membrane (LgBiT-CAAX) or O) to endosomes (FYVE-LgBiT) by 1 µM DAMGO, morphine or PZM21. N,P) Quantification of AUC in M and O, respectively. One-way ANOVA with Tukey’s multiple comparisons, *p<0.05, **p<0.01, ***p<0.001, ****p<0.0001. N=3-4 independent experiments. Error bars represent ± SEM.

To further test this interpretation, we compared the effects of the three MOR agonists in SH-SY5Y cells. Only DAMGO produced a detectable accumulation of active-state MOR on the plasma membrane and endosomes (Figure 5I-L). However, all three agonists clearly increased the level of active-state G_i/o_ both on the plasma membrane and endosomes (Figure 5M-P). The initial level of G protein activity increase produced on the plasma membrane by morphine and PZM21 was lower than that produced by DAMGO and only the DAMGO-induced increase rapidly desensitized (Figure 5M-N). In contrast, all three agonists increased G_i/o_ activity on endosomes and did so to a comparable level (Figure 5O-P). These data are consistent with our results in HEK293 cells (Figure 4) and provide additional support for the interpretation that MOR increases cognate G protein activity on endosomes irrespective of the presence of activated MOR in the endosome membrane.

## Discussion

Over the past two decades, the cell biological understanding of GPCR signaling has evolved to include a second phase of G protein activity on endosomes. However, the present mechanistic understanding of the endosomal activation phase has been elucidated largely through the study of G_s_-coupled GPCRs. G_i_-coupled GPCRs are also known to promote signaling after internalization (24, 33–35), and recent advances in biosensor development have made it possible to spatiotemporally resolve both active-state G_i_-coupled GPCRs (24) and their cognate G proteins (25, 36) in intact cells. Here we apply this approach to delineate the subcellular landscape of cognate G protein activation triggered by MOR, a G_i_-coupled GPCR prototype. By doing so, we reveal the operation of a distinct and additional cellular mechanism for receptor-triggered elevation of cognate G protein activity on endosomes. Importantly, we demonstrate that this mechanism functions in cells with endogenous expression of both receptor and G protein.

Numerous Gs-coupled GPCRs increase endosomal G protein activity by mediating a second round of agonist-dependent allosteric coupling to G proteins on the endosome membrane. This model is supported by multiple lines of evidence, including that endosomal G protein-mediated signaling requires receptors to internalize (9–15) and depends on the presence of activated receptors in the endosome membrane (16–20). There is particularly strong support for this model for receptors that form a stable complex containing receptor, active-state G_s_ and β-arrestin in endosomes (19, 37, 38). Here we show that MOR triggers an elevation of active-state G_i/o_-subclass G proteins on the endosome membrane, but that neither MOR internalization nor MOR activation in endosomes is required. This is not compatible with the current G_s_-coupled GPCR paradigm and reveals the operation of a fundamentally different cellular mechanism for elevating endosomal G protein activity by a G_i_-coupled GPCR. In this second mechanism, we propose that allosteric coupling on the plasma membrane is sufficient to drive an elevation of G protein activity on endosomes without requiring additional allosteric coupling at the endosome. Essentially, the proposed model utilizes trafficking of the activated G protein, rather than of its activating GPCR, to communicate ligand-dependent activation to endosomes (Figure 6).

**Figure 6.**
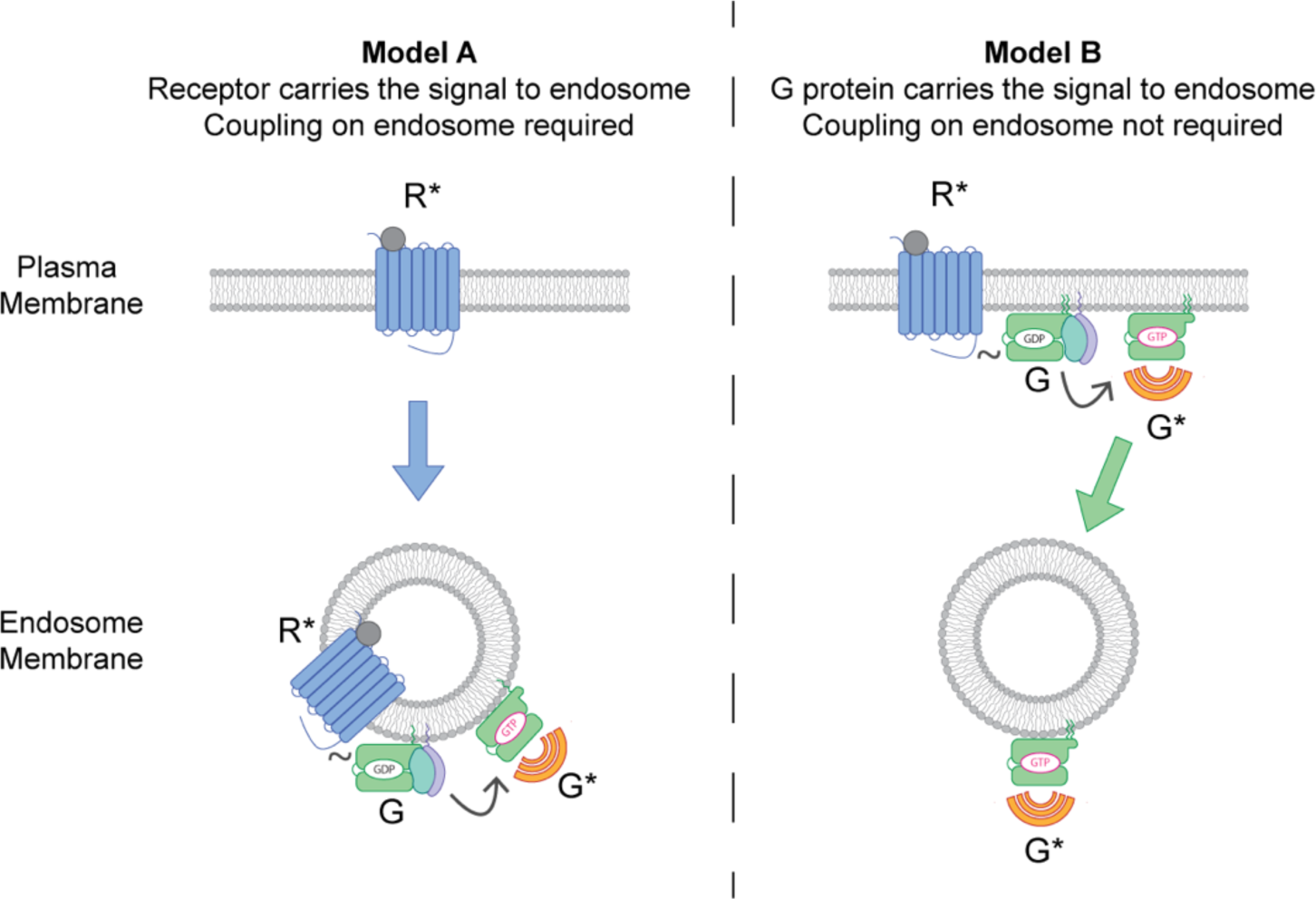
Two distinct models of endosomal G protein activation by GPCRs. In model A, endosomal G protein activity is increased by trafficking of the activated receptor (R*) to endosomes followed by allosteric coupling (∼) on the endosome membrane to generate active-conformation G protein (G*). This model is supported by experimental evidence from G_s_-coupled GPCRs. In model B, supported by the present study, active-conformation G protein (G*) arrives on the endosome membrane following allosteric coupling (∼) with active receptor (R*) on the plasma membrane. These two models differ fundamentally according to 1) whether information is “carried” to endosomes by the GPCR or G protein and 2) the location where allosteric coupling is required.

The present results, by revealing a distinct cellular mechanism of endosomal G protein activation, fundamentally expand our understanding. As our study is limited to MOR, it remains to be determined if this mechanism applies to other G_i_-coupled GPCRs or more broadly to other receptor and G protein subclasses. In the latter regard, we note that G_q_-coupled GPCRs increase G protein activity on endosomes by a mechanism that is dependent on both receptor internalization and the presence of activated receptors in the endosome membrane, consistent with the G_s_-coupled GPCR paradigm Interestingly, a small amount of endosomal G_q_ activity remained when receptor internalization was suppressed, suggesting that this dependence may not be absolute (39). The present results are unique in demonstrating a cellular mechanism of endosomal G protein activation that is dependent solely on receptor activation in the plasma membrane.

Our results strongly support the ability of MOR to increase endosomal G protein activity without requiring receptor activation in the endosome membrane, but they do not rule out the possibility that MOR directly activates G_i/o_ in the endosome membrane to some degree or under some conditions. We observe the presence of activated MOR in the endosome membrane, as reported previously (24), and we now demonstrate this with endogenously expressed receptors. We note that direct activation of endosomal delta opioid receptors has been shown to promote analgesia (34), supporting the importance of receptor activation in endosomes. In addition to increasing G protein activity on endosomes, there is evidence for endosomal GPCR signaling via G protein independent mechanisms (40, 41). Whether endosomal opioid receptors produce physiological effects via G protein dependent or independent mechanisms remains an open question.

Our results raise several additional questions for future investigation. Foremost among these, what is the mechanistic basis for active-state G_i/o_ accumulation on endosomes? Heterotrimeric G proteins are well known to dynamically redistribute in cells (42–44), but it remains unclear how this produces active-state G proteins on endosomes in the absence of local activation by a GPCR. Our antagonist reversal experiments indicate that the active-state lifetime of G_i/o_ is longer on endosomes than on the plasma membrane, as reversal of the G_i/o_ activation signal occurred more slowly on endosomes than the plasma membrane after application of either antagonist (Figure 3E,G). We anticipate that this difference likely involves location-specific differences in regulator of G protein signaling (RGS) proteins that accelerate G_i/o_ inactivation. However, the presently estimated active-state lifetime on endosomes exceeds that measured using purified G proteins in the absence of any RGS protein activity (45). This suggests the existence of additional differences in the biochemical environment of G proteins on endomembranes that enable them to persist in an active conformational state during trafficking, or to be reactivated in transit, but how this occurs is presently unknown.

A second, broader question regards the functional consequences of the MOR-mediated G_i/o_ activity on endosomes. Previous studies indicate that endocytic inhibitors suppress cellular cAMP regulation by G_i_-coupled GPCRs, including opioid receptors, implying the presence of G_i/o_-sensitive adenylyl cyclase(s) on the endosome membrane (24, 35). G_i/o_-subclass G proteins can also regulate many other effectors, in addition to adenylyl cyclase, reviewed in (46). We anticipate that identifying specific G_i/o_-sensitive effectors on endosomes will contribute to a more comprehensive understanding of GPCR signaling selectivity.

In closing, we propose a second cellular mechanism mediating endosomal G protein activation triggered by G_i_-coupled GPCRs. As G_i_-coupled GPCRs comprise the largest GPCR subfamily in mammals (25, 47), we anticipate that these findings will prove broadly relevant to understanding GPCR physiology and help to inform the future development of location-biased therapeutics.

## Materials and Methods

### Cell culture

HEK293 (ATCC CRL-1573) and Gi-KO HEK239A cells (22) were cultured in DMEM (Gibco 1196511) supplemented with 10% FBS (UCSF Cell Culture Facility). Gi-KO HEK293A were grown and plated on dishes coated with collagen (Corning 354236). SH-SY5Y cells were cultured in DMEM:F12 (Gibco 11320082) supplemented with 10% FBS.

### Drugs

Chemicals were purchased from commercial sources as follows: DAMGO (Sigma-Aldrich, E7384), morphine sulfate (Sigma-Aldrich, 1448005), naloxone (Tocris, 05-991-00), CTOP (Tocris, 1578), compound 101 (HelloBio, H2840), pertussis toxin (Sigma-Aldrich, P7208). PZM21 was generously provided by Aashish Manglik (UCSF).

### cDNA constructs

Constructs for pCAGGS-SE-SSF-MOR (24), LgBiT-CAAX (48), and FYVE-LgBiT (48) were previously published. DNA encoding human Gα_i/o/z_ subunits with the internal linker SGGGGTRSGTGGGGS containing restriction sites for KpnI and MluI were synthesized by Twist Bio and cloned into pCAGGS-SE at the KpnI site (site destroyed) by In Fusion cloning (Takara 638946). The insertion sites for each subtype can be found in Table S1. DNA encoding mNeonGreen (Twist Bio) or LgBiT (amplified from LgBiT-CAAX) was further subcloned at the KpnI and MluI sites by In Fusion to create pCAGGS-SE-Gα-mNeonGreen constructs and pCAGGS-SE-G_i1_-LgBiT, respectively. pcDNA-SSF-MOR-LgBiT was constructed by In Fusion insertion of a fragment containing MOR and LgBiT separated by a flexible linker into pcDNA-SSF-MOR digested with BamHI and XbaI. Mitochondria-target LgBiT was constructed by fusing the first 35 amino acids of human Tom20 plus a flexible linker upstream of LgBiT (synthesized by Twist Bio). FYVE-mApple-SmBiT was constructed by inserting SmBiT into pmApple-N1 at the C-terminus of mApple using the Q5 site directed mutagenesis kit (NEB M0492 and M0054) and then inserting the FYVE domain from FYVE-LgBiT at the N-terminus of mApple. pcDNA3-FYVE-mApple was formed by amplifying FYVE-mApple from FYVE-mApple-SmBiT and moving to pcDNA3 at the BamHI site. Nb33-SmBiT and Gi-ED-SmBiT were formed by substituting Nb33 or Gi-ED for miniGs in pcDNA3.1-miniGs-SmBiT (15) at BamHI/HindIII sites. Nb33 was amplified from Nb33-EGFP (24) while Gi-ED was synthesized by Twist Bio. Gi-ED is the first 442 amino acids of human RAP1GAP1 with residues S437, S439, and S441 mutated to alanine as described in (25).

### Live cell confocal microscopy

Image series were acquired using a Nikon Ti inverted microscope controlled by NIS Elements HC v.5.21.03 (Nikon) and fitted with a CSU-22 spinning disk unit (Yokogawa), custom laser launch (100 mW at 405, 488, 561 and 640 nm, Coherent OBIS), Sutter emission filter wheel, Photometrics Evolve Delta EMCCD camera and a temperature- and humidity-controlled chamber (Okolab).

Gi-KO HEK293A cells were plated on collagen-coated 35mM glass-bottom dishes (Cellvis) and transfected 24 hours prior to imaging with 1 µg SSF-MOR, 500 ng Gα-mNeonGreen and 200 ng FYVE-mApple. To surface label SSF-MOR, cells were incubated with anti-FLAG M1 antibody (Millipore Sigma F3040) labeled with Alexa Fluor 647 (Thermo Fisher A20186) for 10 minutes at 37°C and 5% CO_2_. Cells were imaged in imaging media (DMEM, no Phenol Red, 30 mM HEPES, pH 7.4) at 37°C using an Apo TIRF ×100/1.49 numerical aperture oil objective (Nikon). Images were acquired at a 20s frame rate over a 22-minute period with 10 µM DAMGO added at 2 minutes. Images for figures were processed using FIJI software.

### NanoBiT luciferase complementation assays

Cells grown in 6-well dishes were transfected with the appropriate constructs 24 hours before the experiment using Lipofectamine 2000. For HEK293 and Gi-KO HEK293A cells, the amount transfected was as follows: 100 ng of CAAX-LgBiT, FYVE-LgBiT or FYVE-mApple-SmBiT, 500 ng of Nb33-SmBiT or Gi-ED-SmBiT, 250 ng SSF-MOR, and 250 ng tagged Gα constructs where appropriate. For SH-SY5Y cells, the amount of LgBiT constructs was increased to 200 ng and SmBiT constructs to 1 µg. On the day of the assay, cells were lifted using TrypLE (Gibco 12604021) and centrifuged at 500 x g for 3 minutes. Cells were resuspended in assay buffer (in mM: 135 NaCl, 5 KCl, 20 HEPES, 5 D-glucose, 1.8 CaCl2, 0.4 MgCl2, pH 7.4) supplemented with 5 µM coelenterazine-H (Research Products International) to a concentration of 1 x 10^6^. 100 µL of cells were plated into untreated white 96-well plates (Corning 3912). In experiments with PTX, cells were treated overnight with 100 ng/mL PTX prior to assay set up. In experiments with Compound 101 pretreatment, the assay buffer used to plate cells and drug solutions contained 30 µM Compound 101 or DMSO. Total pretreatment time for Compound101 was 20 minutes. Receptor ligands were diluted in assay buffer with 5 µM coelenterazine-H and 50 µL was added during the experiment.

Prior to the start of the assay, the plate was equilibrated for 10 minutes within a platereader warmed to 37°C. Experiments in Figure 2 were performed on a Synergy H4 platereader (Gen5 v.2.05, Biotek). All other experiments were performed on a Tecan Spark platereader. Luminescence values were acquired at 1-minute intervals with an integration time of 500 ms. After a 5-minute baseline, vehicle or drug was added and luminescence was read for an additional 30 minutes. For each well, luminescence was first normalized to the average luminescence during the baseline period. The average luminescence for vehicle treated wells was then subtracted from the drug treated wells at each time point. Each condition was performed as 2-3 technical replicates, which were subsequently averaged. AUC was calculated as the sum of all averaged data points following the drug add.

### Statistical analysis

All data points represent the mean +/− SEM from at least three independent experiments. Statistical tests noted in the figure legends were performed using Prism (Graphpad, v10).

## Acknowledgments

We thank A. Inoue for valuable discussions and for generously sharing the Gi-KO HEK293A cell line, we thank A. Manglik for sharing PZM21 and we thank A. Inoue, J. Li, B. Wysolmerski and E. Blythe for sharing expression constructs. We thank all members of the von Zastrow laboratory for valuable discussion and feedback on the manuscript. Imaging experiments were carried out in the UCSF Center for Advanced Light Microscopy; we thank K. Herrington and S. Kim for their technical support and expertise. This study was supported by research grants from the National Institutes of Health (DA012864, DA010711 and MH120212 to M.v.Z.). N. Fisher was supported by the UCSF IRACDA program (GM081266).

## Author Contributions

N.F. and M.v.Z. designed the research. N.F. performed and analyzed experiments. N.F. and M.v.Z. wrote the paper.

## References

1. D. M. Rosenbaum, S. G. F. Rasmussen, B. K. Kobilka, The structure and function of G-protein-coupled receptors. Nature 459, 356–363 (2009).

2. S. Liu, P. J. Anderson, S. Rajagopal, R. J. Lefkowitz, H. A. Rockman, G protein-coupled receptors: A century of research and discovery. Circ. Res. 135, 174–197 (2024).

3. K. Eichel, M. von Zastrow, Subcellular organization of GPCR signaling. Trends Pharmacol. Sci. 39, 200–208 (2018).

4. C. Kayser, B. Melkes, C. Derieux, A. Bock, Spatiotemporal GPCR signaling illuminated by genetically encoded fluorescent biosensors. Curr. Opin. Pharmacol. 71, 102384 (2023).

5. N. G. Tsvetanova, R. Irannejad, M. von Zastrow, G protein-coupled receptor (GPCR) signaling via heterotrimeric G proteins from endosomes. J. Biol. Chem. 290, 6689–6696 (2015).

6. E. Flores-Espinoza, A. R. B. Thomsen, Beneath the surface: endosomal GPCR signaling. Trends Biochem. Sci. 49, 520–531 (2024).

7. W. Oldham, H. Hamm, Heterotrimeric G protein activation by G-protein-coupled receptors. Nat. Rev. Mol. Cell Biol. 9, 60–71 (2008).

8. W. I. Weis, B. K. Kobilka, The molecular basis of G protein-coupled receptor activation. Annu. Rev. Biochem. 87, 897–919 (2018).

9. S. Ferrandon, et al., Sustained cyclic AMP production by parathyroid hormone receptor endocytosis. Nat. Chem. Biol. 5, 734–742 (2009).

10. D. Calebiro, et al., Persistent cAMP-signals triggered by internalized G-protein-coupled receptors. PLoS Biol. 7, e1000172 (2009).

11. S. J. Kotowski, F. W. Hopf, T. Seif, A. Bonci, M. von Zastrow, Endocytosis promotes rapid dopaminergic signaling. Neuron 71, 278–290 (2011).

12. R. Irannejad, et al., Conformational biosensors reveal GPCR signalling from endosomes. Nature 495, 534–538 (2013).

13. S. Lyga, et al., Persistent cAMP signaling by internalized LH receptors in ovarian follicles. Endocrinology 157, 1613–1621 (2016).

14. J. M. B. Cajulao, E. Hernandez, M. E. von Zastrow, E. L. Sanchez, Glucagon receptor-mediated regulation of gluconeogenic gene transcription is endocytosis-dependent in primary hepatocytes. Mol. Biol. Cell 33, ar90 (2022).

15. E. E. Blythe, M. von Zastrow, β-Arrestin-independent endosomal cAMP signaling by a polypeptide hormone GPCR. Nat. Chem. Biol. 20, 323–332 (2024).

16. T. N. Feinstein, et al., Retromer terminates the generation of cAMP by internalized PTH receptors. Nat. Chem. Biol. 7, 278–284 (2011).

17. T. N. Feinstein, et al., Noncanonical control of vasopressin receptor type 2 signaling by retromer and arrestin. J. Biol. Chem. 288, 27849–27860 (2013).

18. A. Gidon, et al., Endosomal GPCR signaling turned off by negative feedback actions of PKA and v-ATPase. Nat. Chem. Biol. 10, 707–709 (2014).

19. A. R. B. Thomsen, et al., GPCR-G protein-β-arrestin super-complex mediates sustained G protein signaling. Cell 166, 907–919 (2016).

20. X. Tian, et al., The α-Arrestin ARRDC3 Regulates the Endosomal Residence Time and Intracellular Signaling of the β2-Adrenergic Receptor. Journal of Biological Chemistry 291, 14510–14525 (2016).

21. S. K. Gibson, A. G. Gilman, Gialpha and Gbeta subunits both define selectivity of G protein activation by alpha2-adrenergic receptors. Proc. Natl. Acad. Sci. U. S. A. 103, 212–217 (2006).

22. Y. Ono, et al., Generation of Gαi knock-out HEK293 cells illuminates Gαi-coupling diversity of GPCRs. Commun. Biol. 6, 112 (2023).

23. A. K. Finn, J. L. Whistler, Endocytosis of the mu opioid receptor reduces tolerance and a cellular hallmark of opiate withdrawal. Neuron 32, 829–839 (2001).

24. M. Stoeber, et al., A genetically encoded biosensor reveals location bias of opioid drug action. Neuron 98, 963–976.e5 (2018).

25. C. Avet, et al., Effector membrane translocation biosensors reveal G protein and βarrestin coupling profiles of 100 therapeutically relevant GPCRs. Elife 11 (2022).

26. M. Tanowitz, M. von Zastrow, A novel endocytic recycling signal that distinguishes the membrane trafficking of naturally occurring opioid receptors. J. Biol. Chem. 278, 45978–45986 (2003).

27. E. Miess, et al., Multisite phosphorylation is required for sustained interaction with GRKs and arrestins during rapid μ-opioid receptor desensitization. Sci. Signal. 11, eaas9609 (2018).

28. D. E. Keith, et al., Morphine activates opioid receptors without causing their rapid internalization. J. Biol. Chem. 271, 19021–19024 (1996).

29. A. Manglik, et al., Structure-based discovery of opioid analgesics with reduced side effects. Nature 537, 185–190 (2016).

30. S. M. Kazmi, R. K. Mishra, Opioid receptors in human neuroblastoma SH-SY5Y cells: evidence for distinct morphine (mu) and enkephalin (delta) binding sites. Biochem. Biophys. Res. Commun. 137, 813–820 (1986).

31. A. D. Blake, G. Bot, J. C. Freeman, T. Reisine, Differential opioid agonist regulation of the mouse mu opioid receptor. J. Biol. Chem. 272, 782–790 (1997).

32. M. Tanowitz, J. N. Hislop, M. von Zastrow, Alternative splicing determines the post-endocytic sorting fate of G-protein-coupled receptors. J. Biol. Chem. 283, 35614–35621 (2008).

33. F. Mullershausen, et al., Persistent signaling induced by FTY720-phosphate is mediated by internalized S1P1 receptors. Nat. Chem. Biol. 5, 428–434 (2009).

34. N. N. Jimenez-Vargas, et al., Endosomal signaling of delta opioid receptors is an endogenous mechanism and therapeutic target for relief from inflammatory pain. Proc. Natl. Acad. Sci. U. S. A. 117, 15281–15292 (2020).

35. D. S. Eiger, et al., Location bias contributes to functionally selective responses of biased CXCR3 agonists. Nat. Commun. 13, 5846 (2022).

36. M. Maziarz, et al., Revealing the activity of trimeric G-proteins in live cells with a versatile biosensor design. Cell 182, 770–785.e16 (2020).

37. V. L. Wehbi, et al., Noncanonical GPCR signaling arising from a PTH receptor-arrestin-Gβγ complex. Proc. Natl. Acad. Sci. U. S. A. 110, 1530–1535 (2013).

38. A. H. Nguyen, et al., Structure of an endosomal signaling GPCR-G protein-β-arrestin megacomplex. Nat. Struct. Mol. Biol. 26, 1123–1131 (2019).

39. S. C. Wright, et al., BRET-based effector membrane translocation assay monitors GPCR-promoted and endocytosis-mediated Gq activation at early endosomes. Proc. Natl. Acad. Sci. U. S. A. 118, e2025846118 (2021).

40. Y. Daaka, et al., Essential role for G protein-coupled receptor endocytosis in the activation of mitogen-activated protein kinase. J. Biol. Chem. 273, 685–688 (1998).

41. K. A. DeFea, et al., beta-arrestin-dependent endocytosis of proteinase-activated receptor 2 is required for intracellular targeting of activated ERK1/2. J. Cell Biol. 148, 1267–1281 (2000).

42. P. B. Wedegaertner, G protein trafficking. Subcell. Biochem. 63, 193–223 (2012).

43. W. K. Ajith Karunarathne, P. R. O’Neill, P. L. Martinez-Espinosa, V. Kalyanaraman, N. Gautam, All G protein βγ complexes are capable of translocation on receptor activation. Biochem. Biophys. Res. Commun. 421, 605–611 (2012).

44. W. Jang, K. Senarath, G. Feinberg, S. Lu, N. A. Lambert, Visualization of endogenous G proteins on endosomes and other organelles. (2024).

45. M. E. Linder, D. A. Ewald, R. J. Miller, A. G. Gilman, Purification and characterization of Go alpha and three types of Gi alpha after expression in Escherichia coli. J. Biol. Chem. 265, 8243–8251 (1990).

46. V. Syrovatkina, K. O. Alegre, R. Dey, X.-Y. Huang, Regulation, signaling, and physiological functions of G-proteins. J. Mol. Biol. 428, 3850–3868 (2016).

47. A. Inoue, et al., Illuminating G-protein-coupling selectivity of GPCRs. Cell 177, 1933–1947.e25 (2019).

48. Z. Xu, et al., Structural basis of sphingosine-1-phosphate receptor 1 activation and biased agonism. Nat. Chem. Biol. 18, 281–288 (2022).

