## Supplemental information for "Opioid receptors reveal a discrete cellular mechanism of endosomal G protein activation"

**Author Contributions:** N.F. and M.v.Z. designed the research. N.F. performed and analyzed experiments. N.F. and M.v.Z. wrote the paper.

**Competing Interest Statement:** The authors declare no competing interests.

**This PDF file includes:**

Supplemental Figures 1 to 3
Supplemental Table 1

### Figures

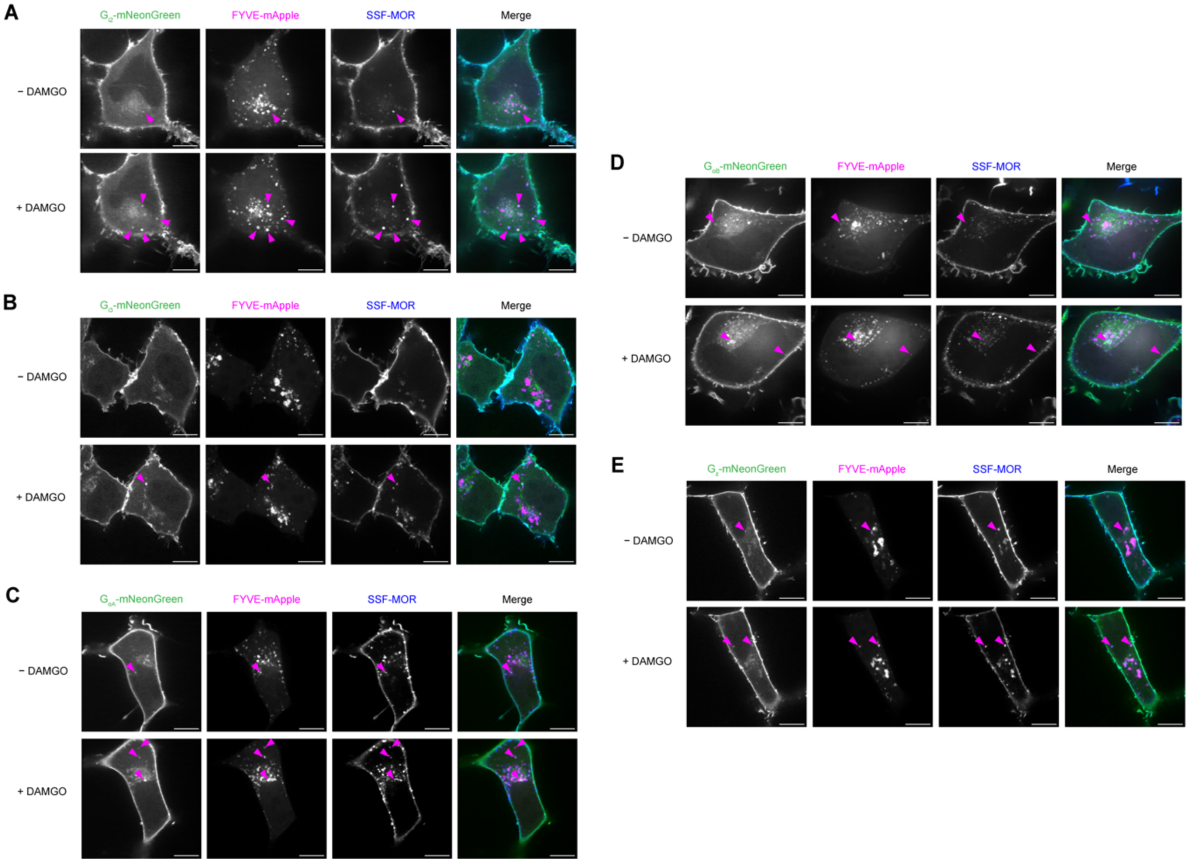


**Figure S1. Localization of other $G_{i/o}$-subclass G proteins upon activation by MOR.** Live imaging by confocal microscopy of Gi-KO HEK293A cells expressing $G_{i2}$-mNeonGreen (A), $G_{i3}$-mNeonGreen (B), $G_{oA}$-mNeonGreen (C), $G_{oB}$-mNeonGreen (D), or $G_z$-mNeonGreen (E) along with the endosome marker FYVE-mApple and SSF-MOR labeled with α-FLAG M1-647. Images depict frames before and after treatment with 10 μM DAMGO for 10-20 minutes. Magenta arrows indicate endosomes positive for Gα-mNeonGreen. Representative images from 2 independent experiments. Scale bar = 10 μm.

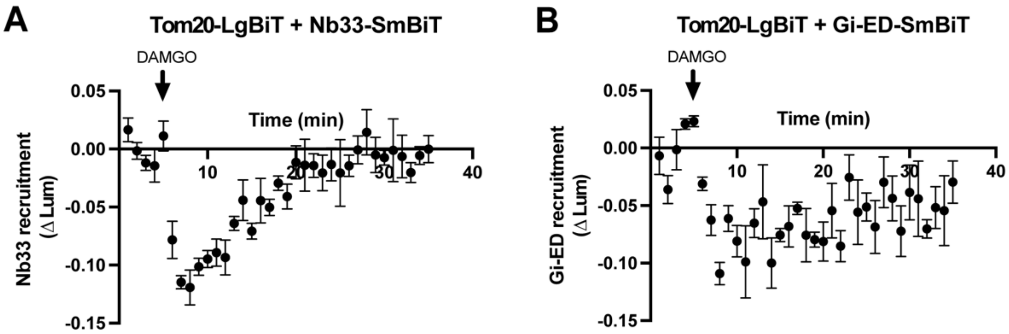


**Figure S2. Nb33-SmBiT and Gi-ED-SmBiT are not recruited to mitochondria following MOR activation.** A) Nb33-SmBiT recruitment to the outer mitochondrial membrane labeled with Tom20-LgBiT following addition of 1 μM DAMGO. B) Gi-ED-SmBiT recruitment to Tom20-LgBiT following addition of 1 μM DAMGO. N=3 independent experiments. Error bars represent ± SEM.

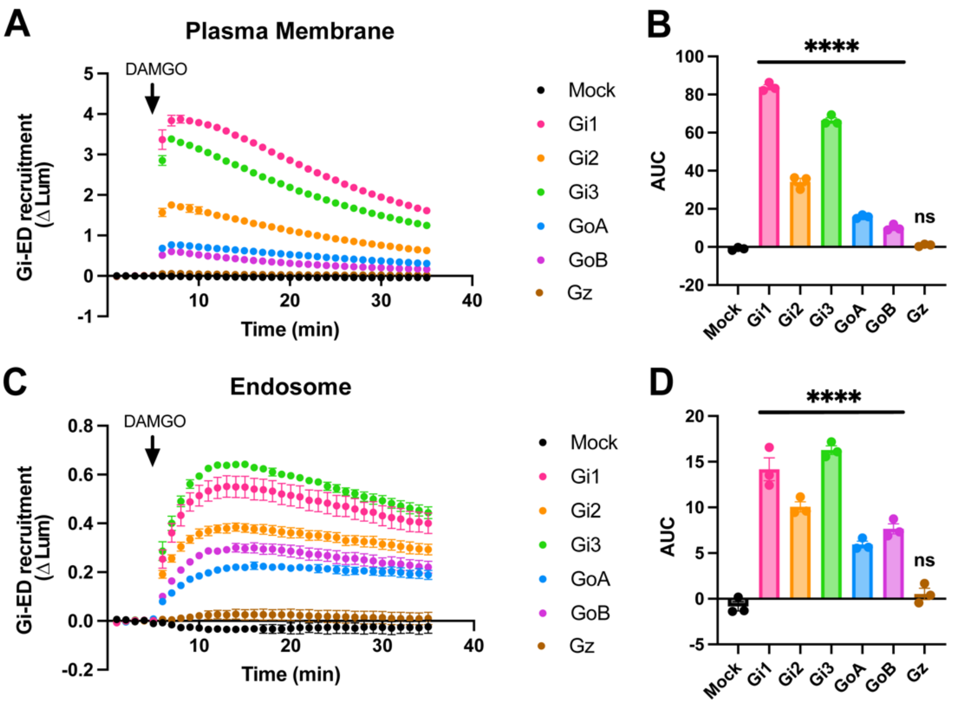


**Figure S3. Activation of $G_{i/o}$ subtypes on the plasma membrane and on endosomes.** A) Gi-ED-SmBiT recruitment to plasma membrane (LgBiT-CAAX) following addition of 1 µM DAMGO in Gi-KO HEK293A cells with re-expression of Gα-mNeonGreen for each $G_{i/o/z}$ subtype. B) Quantification of area under the curve (AUC) for data presented in panel A. One-way ANOVA with Dunnett's multiple comparisons to Mock condition. ****p<0.0001. C) Gi-ED-SmBiT recruitment to endosomes (FYVE-LgBiT) following addition of 1 µM DAMGO in Gi-KO HEK293A cells with re-expression of Gα-mNeonGreen for each $G\alpha_{i/o/z}$ subtype. B) Quantification of area under the curve (AUC) for data presented in panel C. One-way ANOVA with Dunnett's multiple comparisons to Mock condition, ****p<0.0001. N=3 independent experiments. Error bars represent ± SEM.

**Table S1.** Insertion site for mNeonGreen into Gα subtypes

| G protein | Insertion site (amino acids) |
|---|---|
| $G_{i1}$ | 121/122 |
| $G_{i2}$ | 112/115 |
| $G_{i3}$ | 114/115 |
| $G_{oA}$ | 91/92 |
| $G_{oB}$ | 91/92 |
| $G_z$ | 114/115 |
